# A decade of hidden phytoplasmas unveiled through citizen science

**DOI:** 10.1101/2023.01.19.524422

**Authors:** Anne-Sophie Brochu, Antoine Dionne, Mamadou Lamine Fall, Edel Pérez-López

**Affiliations:** Départment de phytologie, Faculté des sciences de l’agriculture et de l’alimentation, Université Laval, Quebec City, Quebec, Canada; Centre de recherche et d’innovation sur les végétaux (CRIV), Université Laval, Quebec City, Quebec, Canada; Institute de Biologie Intégrative et des Systèmes (IBIS), Université Laval, Quebec City, Quebec, Canada; Laboratoire d’expertise et de diagnostic en phytoprotection, MAPAQ, Quebec City, Quebec, Canada; Saint-Jean-sur-Richelieu Research and Development Centre, AAFC, Saint-Jean-sur-Richelieu, Quebec, Canada

**Keywords:** Citizen science, climate change, phytoplasmas, insect vectors, pest monitoring

## Abstract

Climate change is impacting agriculture in many ways, and a contribution from all is required to reduce the imminent loses related to it. Recently, it has been showed that citizen science could be a way to trace the impact of climate change. However, how can citizen science be applied in plant pathology? Here, using as an example a decade of phytoplasma-related diseases reported by growers, agronomists, citizens in general, and confirmed by a government laboratory, we explore a new way of valuing plant pathogens monitoring data deriving from land-users or stakeholders. Through this collaboration we found that in the last decade thirty-four hosts have been affected by phytoplasmas, nine, thirteen and five of these plants were, for the first time, reported phytoplasma hosts in Eastern Canada, in Canada and worldwide, respectively. Another finding of great impact is the first report of a ‘*Ca*. P. phoenicium’-related strain in Canada, while ‘*Ca*. P. pruni’ and ‘*Ca*. P. pyri’ was reported for the first time in Eastern Canada. These findings will have a great impact in the management of phytoplasmas and their insect vectors. Using these insect-vectored bacterial pathogens, we show the needs of new strategies that allow a fast and accurate communication between concerned citizens and those institutions confirming their observations.

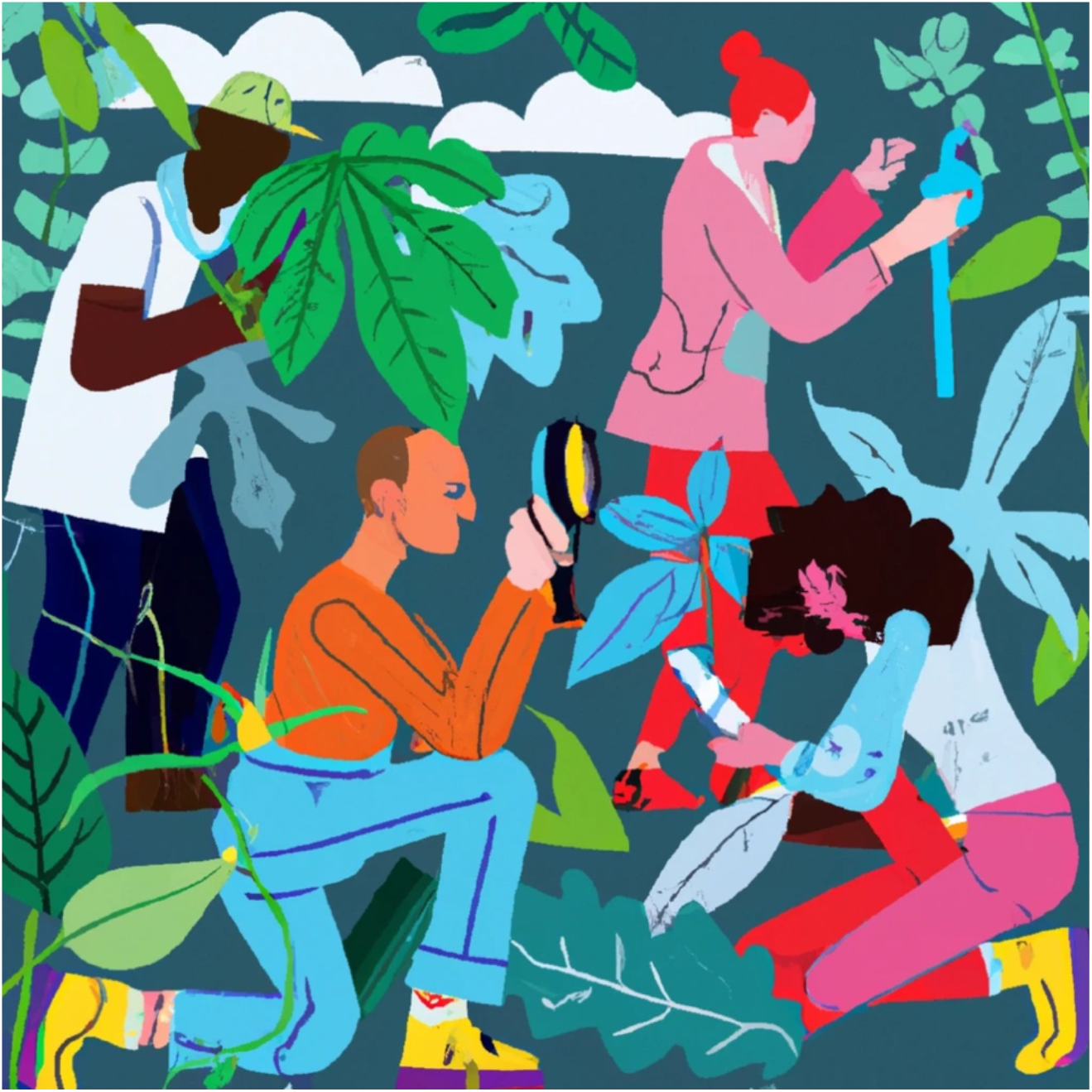

---

Phytoplasmas (‘*Candidatus* Phytoplasma’), are phloem-limited and insect transmitted pathogens associated with a vast number of plant diseases affecting many commercially important crops (Kumari et al. 2019; Pusz-Bochenska et al. 2022). In the last decade, countries like Brazil, India, and China have reported hundreds of new hosts affected by phytoplasmas, evidence of the risk that these pathogens represent in tropical and warm regions (Canale et al. 2020; Rao 2021; Wang et al. 2022). In the coming years, a warmer climate is expected to affect most regions as a consequence of climate change, which will have a serious impact on insect pest distribution, disease incidence, and food security (Ristaino et al. 2021; Outhwaite et al. 2022;). Increases of temperatures can (*i*) accelerate insect’s metabolic rate and herbivory (Dillon et al. 2010; Deutsch et al. 2018), (*ii*) increase the number of insect generations per year (Deutsch et al. 2008), (*iii*) increase the multiplication of phytoplasmas in plants, and the dissemination of the pathogen by insect vectors (Maggi et al. 2014; Bahar et al. 2018; Sabato et al. 2020), and (*iv*) favor the colonization of new niches by expanding their thermal limits, especially for those in temperate regions (Harvey et al. 2020). However, to trace the direct impact of such phenomena is difficult and requires long term studies and active surveillance (Brown et al. 2020).

Recently, the role of citizen science on tracking the effects of climate change has been extensively explored, with the platform *i*Naturalist (https://www.inaturalist.org/) as a successful example, allowing to understand changes in global diversity (Callaghan et al. 2022; Wolf et al. 2022; Shumskaya et al. 2023). However, how could citizen science contribute to identify the incidence of plant diseases during long periods of time? A tool like *i*Naturalist could help to report any encounter of plants affected by some diseases, but for microorganisms like phytoplasmas, a laboratory confirmation is required to ensure that the symptoms observed are related to the presence of the pathogen.

Plant pathology diagnostic laboratories or clinics can play a key role on this kind of studies confirming observations made by growers, agronomists, or just concerned citizens worried by their ornamental plants, community gardens or parks health status. Unfortunately, very often, important information like pathogen diversity or host affected by different disease obtained by those labs remain nondisclosed making difficult to develop efficient evidence-based management strategies, although policy makers are usually informed (Debber et al. 2019). Here, in collaboration with the plant protection and diagnosis laboratory (LEDP, from French *Laboratoire d’expertise et de diagnostic en phytoprotection*) from the Ministry of Agriculture, Fisheries and Food of Quebec, Canada, we are exploring how to make the most of passive surveillance using as a model phytoplasma-related disease reports from the last ten years.

As a standard procedure, once the samples arrive to the LEDP, total DNA is extracted using a CTAB-based method for samples analyzed before 2016, while DNeasy Plant Pro Kit (QIAGEN, CAD) after, and used as template for PCR amplification of the 16S rRNA-encoding gene using phytoplasma universal primers R16F2n/R16R2 as previously described (Gundersen and Lee 1996). Amplicons are later directly sequenced using the amplification primers to confirm the presence of phytoplasma and the results were provided to the client with a yes or no answer to the question if the sample is indeed infected by the pathogen. As part of our study, in addition to information regarding host and year of collection, we also had access to the forward and reverse 16S rRNA-encoding gene sequence amplified from the samples. To identify the phytoplasma species, the sequences were assembled using the Staden package (Bonfield and Whitwham 2010) and compared with references from GenBank using BLAST (http://www.ncbi.nlm.nih.gov). Phylogenetic analysis was conducted using the Neighbor Joining algorithm in MEGA X (Kumar et al. 2018), and bootstrapping 1000 times to estimate stability. *Acholeplasma laidlawii* strain PG-8 A (U14905) was used as outgroup to root the tree.

In terms of phytoplasma diversity, the vast majority of the phytoplasma strains identified in the last decade affecting plants in East Canada are ‘*Candidatus* Phytoplasma asteris’-related strains (Fig. 1A, Table 1). Interestingly, we found, for the first time in Canada, the presence of ‘*Ca*. P. phoenicium’-related strains (16SrIX), and for the first time in Quebec and East Canada, the presence of ‘*Ca*. P. pruni’-related strains (16SrIII) and ‘*Ca*. P. pyri’-related strains (16SrX) (Fig. 1A, Table 1). Here we found a ‘*Ca*. P. phoenicium’-related strain infecting blueberry plants, which previous studies showed to be typically infected by ‘*Ca*. P. asteris’ causing blueberry bushy stunt disease (BbSP) (Pérez-López et al. 2019; Arocha-Rosete et al. 2019; Hammond et al. 2021). However, blueberry plants have been reported infected by ‘*Ca*. P. phoenicium’ in New Jersey, USA, also causing BbSP in 2013 (Bagadia et al. 2013), just a year before ‘*Ca*. P. phoenicium’-infected sample was collected in Quebec (Table 1). In Canada, ‘*Ca*. P. pruni’ was previously reported affecting peach trees, milkweed, clover, and chokecherry in Ontario (Davis et al. 1990; Gundersen et al. 1996; Lee et al. 1993; Wang and Hiruki 2005), chokecherry in Saskatchewan (Wang and Hiruki 2005), and pin cherry in Alberta (Wang and Hiruki 2005), while now we report that in 2012 a blackcurrant tree was positive for this phytoplasma species (Fig. 1A, Table 1). Similarly, ‘*Ca*. P. pyri’ was previously reported in Canada but only inffecting pear trees in British Columbia and Ontario (Seemüller and Schneider 2004; Hunter et al. 2010), same host affected by this phytoplasma species in Quebec in 2016 and later in 2019 (Fig. 1A, Table 1).

**Fig. 1.**
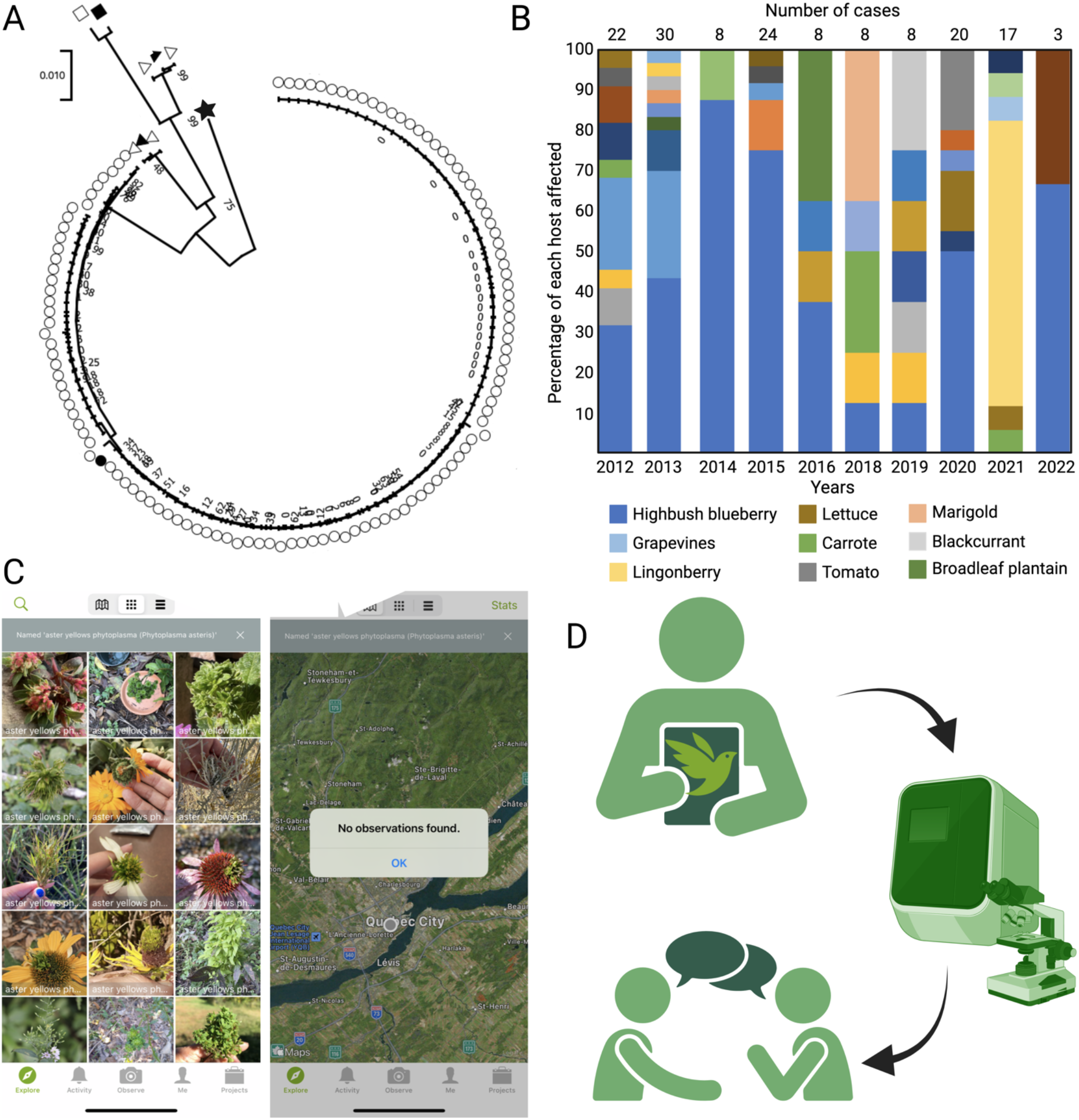
Phylogenetic analysis of 16S sequences generated by the LEDP in the last decade (**A**). Phylogenetic analysis was performed using the Neighbor Joining algorithm using 1000 replicates, as described in the main text. Circles are signaling ‘*Ca*. P. asteris’-related strains, square is marking ‘*Ca*. P. phoenicium’-related strain, triangle to the right is marking ‘*Ca*. P. pruni’-related strains, and triangle to the left is marking ‘*Ca*. P. pyri’-related strains. Filled labels represent reference species and star is the outgroup. Representation of the host diversity e incidence of phytoplasmas in the last decade (**B**). *i*Naturalist search for ‘aster yellow phytoplasma’ shows several results in different geographic locations but no results in Quebec despite knowing that hundreds of cases have been confirmed (**C**). Schematic representation of the model proposed by us to increase the impact of citizen science (**D**).

**Table 1.** Full information of those samples affected by phytoplasmas identified and analyzed in Quebec from 2012 to 2022.

| Host | No. samples<br>(year) | ‘*Candidatus*<br>Phytoplasma’<br>species | Genbank<br>accession no. | New host |
| --- | --- | --- | --- | --- |
| Highbush Blueberry<br>(*Vaccinium*<br>*corymbosum*) | 6 (2012), 12<br>(2013), 6 (2014),<br>18 (2015), 2<br>(2016), 1 (2018),<br>1 (2019), 10<br>(2020), 1 (2022) | ‘*Candidatus*<br>Phytoplasma asteris’ | OQ211246-OQ211266,<br>OQ211268-OQ211303 | No |
|  | 1 (2014) | ‘*Candidatus*<br>Phytoplasma<br>phoenicium’ | OQ211267 | Canada |
| Grapevines (*Vitis*<br>*vinifera*) | 5 (2012), 8<br>(2013), 1 (2015) | ‘*Candidatus*<br>Phytoplasma asteris’ | OQ211335-OQ211348 | No |
| Lingonberry<br>(*Vaccinium vitis-*<br>*idaea*) | 12 (2021) | ‘*Candidatus*<br>Phytoplasma asteris’ | OQ211466-OQ211477 | No |
| Lettuce (*Lactuca*<br>*sativa*) | 1 (2012), 3<br>(2020), 1 (2021) | ‘*Candidatus*<br>Phytoplasma asteris’ | OQ214210-OQ214214 | Eastern<br>Canada |
| Tomato (*Solanum*<br>*lycopersicum*) | 3 (2020) | ‘*Candidatus*<br>Phytoplasma asteris’ | OQ214878-OQ214880 | Canada |
| Carrot<br>(*Daucus carota*) | 1 (2012), 2<br>(2018), 1 (2021) | ‘*Candidatus*<br>Phytoplasma asteris’ | OQ214892-OQ214895 | No |
| Raspberry (*Rubus*<br>*fruticosus*) | 1 (2012), 1<br>(2018), 1 (2019) | ‘*Candidatus*<br>Phytoplasma asteris’ | OQ214897-OQ214899 | Canada |
| Lowbush Blueberry<br>(*Vaccinium*<br>*angustifolium*) | 3 (2015) | ‘*Candidatus*<br>Phytoplasma asteris’ | OQ214925-OQ214927 | No |
| Marigold (*Tagetes patula*) | 3 (2018) | ‘*Candidatus*<br>Phytoplasma asteris’ | OQ214954-OQ214956 | Eastern<br>Canada |
| Elderberry (*Sambucus nigra*) | 2 (2012), 1 (2020) | ‘*Candidatus*<br>Phytoplasma asteris’ | OQ214932-OQ214934 | World |
| Apple (*Malus domestica*) | 3 (2013) | ‘*Candidatus*<br>Phytoplasma asteris’ | OQ215319-OQ215321 | Canada |
| Honeysuckle<br>(*Lonicera kamtschatica*) | 1 (2013), 1 (2019) | ‘*Candidatus*<br>Phytoplasma asteris’ | OQ214952-OQ214953 | World |
| Bell pepper<br>(*Capsicum annuum*) | 2 (2019) | ‘*Candidatus*<br>Phytoplasma asteris’ | OQ215312-OQ215313 | Canada |
| Garlic (*Allium sativum*) | 1 (2013), 1 (2020) | ‘*Candidatus*<br>Phytoplasma asteris’ | OQ215314-OQ215315 | Eastern<br>Canada |
| Blackcurrant (*Ribes nigrum*) | 1 (2012) | ‘*Candidatus*<br>Phytoplasma pruni’ | OQ215738 | Canada |
|  | 1 (2012) | ‘*Candidatus*<br>Phytoplasma asteris’ | OQ215739 |  |
| Pear (*Pyrus communis*) | 1 (2016), 1 (2019) | ‘*Candidatus*<br>Phytoplasma pyri’ | OQ215383-OQ215384 | Eastern<br>Canada |
| Speedwells (*Veronica persica*) | 1 (2016), 1 (2020) | ‘*Candidatus*<br>Phytoplasma asteris’ | OQ215735-OQ215736 | Canada |
| Celery (*Apium graveolens*) | 1 (2013) | ‘*Candidatus*<br>Phytoplasma asteris’ | OQ215745 | Eastern<br>Canada |
| Broccoli (*Brassica oleracea*) | 1 (2018) | ‘*Candidatus*<br>Phytoplasma asteris’ | OQ215754 | Canada |
| Lily (*Lilium candidum*) | 1 (2013) | ‘*Candidatus*<br>Phytoplasma asteris’ | OQ215755 | Canada |
| Peony (*Paeonia officinalis*) | 1 (2012) | ‘*Candidatus*<br>Phytoplasma asteris’ | OQ215756 | Canada |
| Larkspur (*Delphinium*<br>sp.) | 1 (2021) | ‘*Candidatus*<br>Phytoplasma asteris’ | OQ215779 | Canada |
| Potato (*Solanum tuberosum*) | 1 (2015) | ‘*Candidatus*<br>Phytoplasma asteris’ | OQ221893 | Eastern<br>Canada |
| Wheat (*Triticum aestivum*) | 1 (2014) | ‘*Candidatus*<br>Phytoplasma asteris’ | OQ215792 | Canada |
| Marshmallow<br>(*Althaea officinalis*) | 1 (2012) | ‘*Candidatus*<br>Phytoplasma asteris’ | OQ215793 | World |
| New Guinea<br>impatiens (*Impatiens hawkeri*) | 1 (2015) | ‘*Candidatus*<br>Phytoplasma asteris’ | OQ215794 | World |
| Soybean (*Glycine max*) | 1 (2019) | ‘*Candidatus*<br>Phytoplasma asteris’ | OQ215796 | Canada |
| Bonesets (*Eupatorium perfoliatum*) | 1 (2013) | ‘*Candidatus*<br>Phytoplasma asteris’ | OQ216289 | World |
| Broadleaf plantain<br>(*Plantago major*) | 1 (2016) | ‘*Candidatus*<br>Phytoplasma asteris’ | OQ216290 | Eastern<br>Canada |
| Gloxinia (*Gloxinia perennis*) | 1 (2013) | ‘*Candidatus*<br>Phytoplasma asteris’ | OQ216489 | Canada |
| Dahlia (*Dahlia pinnata*) | 1 (2022) | ‘*Candidatus*<br>Phytoplasma asteris’ | OQ216557 | Eastern<br>Canada |
| Willow (*Salix alba*) | 1 (2021) | ‘*Candidatus*<br>Phytoplasma asteris’ | OQ216558 | Eastern<br>Canada |
| Alfalfa (*Medicago sativa*) | 1 (2020) | ‘*Candidatus*<br>Phytoplasma asteris’ | OQ216560 | Eastern<br>Canada |
| Aster (*Aster* sp.) | 1 (2021) | ‘*Candidatus*<br>Phytoplasma asteris’ | OQ216559 | Eastern<br>Canada |

In terms of hosts affected by phytoplasmas, the results show a broad host range. Overall, we found 34 hosts inffected by these pathogenic bacteria from 2012 to 2022, distributed in a total of 148 samples. For nineteen of those hosts only one sample was analyzed but considering that this number is influenced by several factors like, price of the tests, time availability of the observer/reporter to perform extensive surveys in the field, and economic or ecologic importance of the infected plant, we believe that reporting those hosts is necessary. Interestingly, twenty-eight of those hosts are new reports organized into three categories: (*i*) new phytoplasma host for Eastern Canada, (*ii*) new phytoplasma host for Canada, and (*iii*) new phytoplasma host worldwide. In the first category we have nine hosts: lettuce, celery, alfalfa, pear, peony, garlic, broadleaf plantain, marigold, and aster (Table 1). All these have been previously reported in Canada affected by phytoplasmas (Olivier et al. 2009). For example, celery and garlic have been reported infected by ‘*Ca*. P. asteris’ in Alberta, pear trees, as we mentioned before, have been reported affected by ‘*Ca*. P. pyri’ in British Columbia, and broadleaf plantain has been reported affected by ‘*Ca*. P. asteris’ in Manitoba (Olivier et al. 2009) (Fig. 1B, Table 1). In the second category, new hosts for Canada, we have thirteen including blackcurrant, raspberry, peony, apple trees, lily, gloxinia, wheat, soy, tomatoes, speedwells, broccoli, pepper, and larkspur (Olivier et al. 2009) (Table 1). For example, blackcurrant has been previously reported affected by ‘*Ca*. P. asteris’ in Czech Republic (ŠPak et al. 2004), raspberry affected by ‘*Ca*. P. hyspanicum’-related strains in Mexico (Pérez-López et al. 2017), lily has been reported infected by ‘*Ca*. P. asteris’-related strains in Mexico (Cortés-Martínez et al. 2013), and peonies by ‘*Ca*. P. solani’-related strains in China (Gao et al. 2012). An interesting finding is that apple trees were found positive for phytoplasma in 2013 in Quebec, same year when there was a big controversy around apple trees that could be affected by the quarantine pest apple proliferation phytoplasma (‘*Ca*. P. mali’), but fortunately in Quebec was ‘*Ca*. P. asteris’ (Table 1). The last category includes those hosts that haven’t been reported to be infected by phytoplasma before at the global scale, including elderberry, honeysuckle, marshmallow, New Guinea impatiens, and bonesets (Table 1). After an extensive review of literature, we were not able to find any report of these plant species infected by phytoplasmas elsewhere before.

Another element that we can discuss is the prevalence of phytoplasma-related disease, as well as the diversity of hosts affected every year (Fig. 1B). Generally, we observed that those years with high incidence have also a high diversity of infected hosts, except for 2015, when was experienced a high number of cases, but almost exclusively limited to blueberry plants infected by BbSP (Fig. 2B). Although without concrete evidence, we believe that many growers and agronomists could send samples to do confirmation during one or two growing seasons and after they will be able to recognize the symptoms and proceed to eliminate symptomatic plants to avoid spreading of the disease. This could explain why we see how the number of samples for certain hosts is high in some years and then decreases in the following years. That could be the case for BbSP, a disease we know has been present during all the past decade in Quebec (Pérez-López et al. 2019; Rosete et al. 2019; Hammond et al. 2021), but we only see a high number of samples up to 2015, and an increase later in 2020 (Table 1, Fig. 1B). Based on these observations, we should be cautious correlating the number of cases analyzed by the LEDP with the real incidence of certain phytoplasma-related disease.

Through this study we are highlighting the role of citizen science and plant pathology laboratories on following the incidence of plant diseases. To this day, we don’t have an open platform that could be used to record in real time symptoms or the presence of certain diseases. *i*Naturalist has been used also for this purpose and we were able to find aster yellow phytoplasma reports in several locations, although we didn’t find any report in Quebec (Fig. 1C). United Kingdom, New Zealand, South Africa, and United Sates have been exploring citizen science to study the distribution of plant pathogens and pests for several years (Hulbert 2017; Ryan et al. 2018; Brown et al. 2020; de Groot et al. 2023), with some examples here in Canada, mainly to study the distribution of insect pests (Martel 2020). Currently, there is a citizen project ongoing focused on the plant pathogen *Sclerotinia sclerotiorum* affecting common bean (https://www.pulsebreeding.ca/research/canadian-sclerotinia-initiative) This project follows a model organize in three main steps: incidence and collection performed by citizens, analysis performed by a laboratory to later release results and confirmation of the identity of the pathogen. Nonetheless, how can transparency be increased in the process? We propose the following suggestions to ensure fast availability and quality of the data generated through citizen science in Canada and elsewhere: (1) registration of symptoms in an interactive app or database like *i*Naturalist by observer or analyzer, which would also guarantee to have a photographic archive of symptoms, (2) laboratory confirmation by academic, government, or privet institutions interested on follow up the specific disease or pest, and (3) divulgation of the results through the same app or database, reports or peer-review publications (Fig. 1D). This system is not sustainable for all plant pathogens due to the high volume of analysis, but for diseases expected to be highly influenced by climate change like those transmitted by insect vectors, is an option can be used as early warning to anticipate threats from domestic, latent, emerging, new and transboundary pests and to be ready for possible outbreaks. Through this collaboration between the provincial government and academia, we have been able to highlight an issue that goes further than phytoplasmas and can have an impact in all of us.

## AKNOWLEDGEMENTS

We would like to thank the LEDP from MAPAQ and his staff and NSERC for the funding to EPL through the Discovery grant.

## REFERENCES

Arocha-Rosete, Y., Lambert, L., Joly-Séguin, V., Michelutti, R., Schilder, A., and Bertaccini, A. 2019 Surveys reveal a complex association of phytoplasmas and viruses with the blueberry stunt disease on Canadian blueberry farms. Ann. Appl. Biol. 174:142–152.

Bagadia, P. G., Polashock, J., Bottner-Parker, K. D., Zhao, Y., Davis, R. E. et al. 2013. Characterization and molecular differentiation of 16SrI-E and 16SrIX-E phytoplasmas associated with blueberry stunt disease in New Jersey. Mol. Cell Probes. 27:90–97.

Bahar, M. H., Wist, T., Bekkaoui, D. et al. 2018. Aster leafhopper survival and reproduction, and Aster yellows transmission under static and fluctuating temperatures, using ddPCR for phytoplasma quantification. Sci. Rep. 8: 227.

Bonfield, J. K., and Whitwham, A. 2010. Gap5 - editing the billion fragment sequence assembly. Bioinformatics 26: 1699–1703.

Brown, N., Pérez-Sierra, A., Crow, P., and Parnell, S. 2020. The role of passive surveillance and citizen science in plant health. CABI Agric. Biosci. 1:17.

Callaghan, C. T., Mesaglio, T., Ascher, J. S., Brooks, T. M., Cabras, A. A., Chandler. M., et al. 2022. The benefits of contributing to the citizen science platform iNaturalist as an identifier. PLoS Biol. 20(11): e3001843.

Canale, M. C., and Bedendo, I. P. 2020. Report of ‘Candidatus Phytoplasma hispanicum’ (16SrXIII-E) Associated with Cauliflower Stunt in São Paulo State, Brazil, and Balclutha hebe as Its Potential Vector. Plant Disease. 104:3.

Cortés-Martínez, N. E., Valadez-Moctezuma, E., Zelaya-Molina, L. X., and Marbán-Mendoza, N. 2008. First Report of ‘Candidatus Phytoplasma asteris’-Related Strains Infecting Lily in Mexico. Plant Disease. 92:6.

Crow, P., Perez-Sierra, A., Kavčič, A., Lewthwaite, K., Kolšek, M., Ogris, N., Piškur, B., Veenvliet, J. K., Zidar, S., Sancisi-Frey, S., and de Groot, M. 2020. Using Citizen Science to monitor the spread of tree pests and diseases: outcomes of two projects in Slovenia and the UK. M. Biol. Invasions. 11(4): 703–719.

Davis, R.E., Lee, I.-M., Douglas, S.M., and Dally, E.L. 1990. Molecular cloning and detection of chromosomal and extrachromosomal DNA of the mycoplasmalike organism associated with little leaf disease in periwinkle (Catharanthus roseus). Mol. Plant Pathol. 80:789–793.

de Groot, M., Pocock, M. J. O., Bonte, J., Fernandez-Conradi, P., and Valdés-Correcher, E. 2023. Citizen Science and Monitoring Forest Pests: a Beneficial Alliance? Curr. For. Rep. 9:15–32.

Deutsch, C. A., Tewksbury, J. J., Huey, R. B., Sheldon, K. S., Ghalambor, C. K., Haak, D. C. and Martin, P. R. 2008. Impacts of climate warming on terrestrial ectotherms across latitude. Proc. Natl. Acad. Sci. U.S.A. 105:6668–6672.

Deutsch, C. A., Tewksbury, J. J., Tigchelaar, M., Battisti, D. S., Merrill, S., Huey, R. B., and Naylor, R. L. 2018. Increase in crop losses to insect pests in a warming climate. Science. 361(6405):916–919.

Dillon, M. E., Wang, G., and Huey, R. B. 2010. Global metabolic impacts of recent climate warming. Nature. 467:704–706.

Gao, Y., Qiu, P. P., Liu, W. H., Su, W. M., Gai, S. P., Liang, Y. C., and Zhu, X.P. 2013. Identification of ‘Candidatus Phytoplasma solani’ associated with tree Peony yellows disease in China. J. Phytopathol. 161:197–200.

Gundersen, D. E., and Lee, I. M. 1996. Ultrasensitive detection of phytoplasmas by nested-PCR assays using two universal primer pairs. Phytopathol. Mediterr. 35:144–151.

Gundersen, D.E., Lee, I.-M., Schaff, D.A., Harrison, N.A., Chang, C.J., Davis, R.E., and Kingsbury, D.T. 1996. Genomic diversity and differentiation among phytoplasma strains in 16S rRNA groups I (aster yellows and related phytoplasmas) and III (X-disease and related phytoplasmas). Int. J. Bacteriol. 46:64–75.

Hammond, C., Pérez-López, E., Town, J. et al. 2021. Detection of blueberry stunt phytoplasma in Eastern Canada using cpn60-based molecular diagnostic assays. Sci. Rep. 11:22118.

Harvey, J. A., Heinen, R., Gols, R., and Thakur, M. P. 2020. Climate change-mediated temperature extremes and insects: From outbreaks to breakdowns. Glob Chang Biol. 26:6685–6701.

Hulbert, J. 2017. “Pathogen hunters”: citizen scientists track plant diseases to save species. The Conversation. https://theconversation.com/amp/pathogen-hunters-citizen-scientists-track-plant-diseases-to-save-species-73710.

Hunter, D. M., Svircev, A. M., Kaviani, M., Michelutti, R., Wang, L., and Thompson, D. 2010. First Report of Pear Decline Caused by ‘Candidatus Phytoplasma pyri’ in Ontario, Canada. Plant Disease. 94:5.

Kumar, S., Stecher, G., Li, M., Knyaz, C., and Tamura, K. 2018. MEGA X: molecular evolutionary genetics analysis across computing platforms. Molec. Biol. Evol. 35:1547–1549.

Kumari, S., Nagendran, K., Rai, A. B., Singh, B., Rao, G. P. and Bertaccini, A. 2019. Global Status of Phytoplasma Diseases in Vegetable Crops. Front. Microbiol. 10:1349.

Lee, I.-M., Hammond, R.W., Davis, R.E., and Gundersen, D.E. 1993. Universal amplification and analysis of pathogen 16SrDNA for classification and identification of mycoplasmalike organisms. Phytopathol. 83:834–842.

Maggi, F., Galetto, L., Marzachì, C., and Bosco, D. 2014. Temperature-dependent transmission of ‘Candidatus Phytoplasma asteris’ by the vector leafhopper Macrosteles quadripunctulatus Kirschbaum. Entomologia. 2:202.

Martel, V. 2020. Citizen scientist spots a newcomer on Canadian elm trees. Government of Canada. https://www.nrcan.gc.ca/simply-science/citizen-scientist-spots-newcomer-on-canadian-elm-trees/23000.

Olivier, C.Y., Lowery, D.T., and Stobbs, L.W. 2009. Phytoplasma diseases and their relationships with insect and plant hosts in Canadian horticultural and field crops. Can. Entomol. 141:425–462.

Outhwaite, C.L., McCann, P., and Newbold, T. 2022. Agriculture and climate change are reshaping insect biodiversity worldwide. Nature. 605:97–102.

Perez-Lopez, E., Vincent, C., Moreau, D., Hammond, C., Town, J., and Dumonceaux, T. J. 2019. A novel ‘Candidatus Phytoplasma asteris’ subgroup 16SrI-(E/AI) AI associated with blueberry stunt disease in eastern Canada. Int. J. Syst. Evol. Microbiol. 69:322–332.

Pérez-López, E., Rodríguez-Martínez, D., Olivier, C.Y. et al. 2017. Molecular diagnostic assays based on cpn60 UT sequences reveal the geographic distribution of subgroup 16SrXIII-(A/I)I phytoplasma in Mexico. Sci Rep. 7: 950.

Pusz-Bochenska, K., Perez-Lopez, E., Wist, T. J., Bennypaul, H., Sanderson, D., Green, M., and Dumonceaux, T. J. 2022. Multilocus sequence typing of diverse phytoplasmas using hybridization probe-based sequence capture provides high resolution strain differentiation. Front. Microbiol. 13:959562

Rao, G. P. 2021. Our understanding about phytoplasma research scenario in India. Indian Phytopathol. 74:371–401.

Ristaino, J. B., Anderson, P. K., Bebber, D. P., Brauman, K. A., Cunniffe, N. J., Fedoroff, N. V., Finegold, C., Garrett, K. A., Gilligan, C. A., Jones, C. M., et al. 2021. The persistent threat of emerging plant disease pandemics to global food security. Proc. Natl. Acad. Sci. U.S.A. 118: e2022239118.

Ryan, S. F., Adamson, N. L., Aktipis, A., Andersen, L. K., Austin, R., Barnes, L., et al. 2018. The role of citizen science in addressing grand challenges in food and agriculture research. Proc. R. Soc. B. 285: 20181977.

Sabato, E. O., Landau, E. C., Barros, B. A., and Oliveira, C. M. 2020. Differential transmission of phytoplasma and spiroplasma to maize caused by variation in the environmental temperature in Brazil. Eur. J. Plant Pathol. 157(1):163–171.

Seemüller, E., and Schneider, B. 2004. ‘Candidatus Phytoplasma mali’, ‘Candidatus Phytoplasma pyri’ and ‘Candidatus Phytoplasma prunorum’, the causal agents of apple proliferation, pear decline and European stone fruit yellows, respectively. Int. J. Syst. Evol. Microbiol. 54:1217–1226.

Shumskaya, M., Filippova, N., Lorentzen, L. et al. 2023. Citizen science helps in the study of fungal diversity in New Jersey. Sci. Data. 10:10.

ŠPak, J., Pribylová, J., Kubelková, D., and ŠPaková, V. 2004. The Presence of Phytoplasma in Black Currant Infected with the Blackcurrant Reversion Disease. J. Phytopathol. 152 (11-12):600–605.

Wang, K., and Hiruki, C. 2005. Distinction between phytoplasmas at the subgroup level detected by heteroduplex mobility assay. Plant Pathol. 54: 625–633.

Wang, X.-Y., Zhang, R.-Y., Li, J., Li, Y.-H., Shan, H.-L., Li, W.-F., and Huang, Y.-K. 2022. The Diversity, Distribution and Status of Phytoplasma Diseases in China. Front. Sustain. Food Syst. 6:943080.

Wolf, S., Mahecha, M.D., Sabatini, F.M. et al. 2022. Citizen science plant observations encode global trait patterns. Nat. Ecol. Evol. 6:1850–1859.

